# Fi-score: a novel approach to characterise protein topology and aid in drug discovery studies

**DOI:** 10.1101/2020.06.25.171124

**Authors:** Austė Kanapeckaitė, Claudia Beaurivage, Matthew Hancock, Erik Verschueren

## Abstract

Target evaluation is at the centre of rational drug design and biologics development. In order to successfully engineer antibodies, T-cell receptors or small molecules it is necessary to identify and characterise potential binding or contact sites on therapeutically relevant target proteins. Currently, there are numerous challenges in achieving a better docking precision as well as characterising relevant sites. We devised a first-of-its-kind *in silico* protein fingerprinting approach based on dihedral angle and B-factor distribution to probe binding sites and sites of structural importance. In addition, we showed that the entire protein regions or individual structural subsets can be profiled using our derived fi-score based on amino acid dihedral angle and B-factor distribution. We further described a method to assess the structural profile and extract information on sites of importance using machine learning Gaussian mixture models. In combination, these biophysical analytical methods could potentially help to classify and systematically analyse not only targets but also drug candidates that bind to specific sites which would greatly improve pre-screening stage, target selection and drug repurposing efforts in finding other matching targets.

## Introduction

The identification of lead compounds showing pharmacological promise is the focal point of early-stage drug discovery. While large libraries of compounds against a therapeutically-relevant target are subjected to high-throughput screening (HTS) to select new lead compounds, this method becomes more and more supplemented or preceded by in *silico* HTS within the pharmaceutical industry. This shift in the paradigm can be attributed to the high costs and time-consuming nature of the design and completion of HTS screens [1]. In contrast, early stage *in silico* screening offers not only a better understanding of relevant biological topology, potential active sites but also allows progressive optimisation of the pharmacological properties and potency of selected compounds. Yet, structurally complex sites or sites with a wide dynamic range pose a challenge; especially, when selecting between a family of targets or targets with similar topology [2].

While the human genome contains about 25,000 genes, only about 10% of expressed proteins are amenable to small-molecule modulation and less than half of that subset have therapeutic potential. In addition, development of therapeutic compounds have a very low success rate as less than 2% of these lead compounds succeed to get to the market [2]–[4].The picture gets even more complicated for immunotherapeutics development as target compounds can have potentially far reaching side effects and the identification and validation of disease-specific targets is also complicated by the fact that numerous proteins can undergo significant conformational changes throughout their immune cycle [2], [3]. Consequently, knowing the binding sites and physicochemical properties of the target protein prior to screening or optimisation of lead compounds would be extremely beneficial in terms of screening cost reduction and faster turnaround or identification of other potential back-up pharmaceutical candidates.

Most currently marketed small-molecule drugs are developed to target protein–ligand interactions (PLI)[5] and this information is primarily provided by crystallographic analysis. Crystallographic structure analysis has revealed that PLI sites are hydrophobic pockets concave in shape with more complex topological features than those found on protein surfaces, but they can also be relatively flat and large [5]–[9]. As a result, computational analysis to probe potential binding sites of proteins exploit these features to evaluate energetics, cavity geometry and physicochemical properties of a potential binding pocket. However, there are additional challenges as the selected sites might be topologically constrained and because of growing computational costs broader conformational changes may not be incorporated into the binding grid analysis. Furthermore, there is not one universal algorithm developed that could be suitable for all scenarios; therefore, we aimed to combine multiple levels of analysis, capturing B-factor values and dihedral angle structure to establish a comparative measure of physicochemical characteristics of a protein of interest that could be used to analyse a single motif, expanded to a site or the whole protein [10].

Protein dihedral angles contain information on the local protein conformation in such a way that a protein backbone conformation can be highly accurately rebuilt based on the native dihedral angles. Extracting this information can facilitate in narrowing down the conformational space, which in turn can be superimposed on specific physicochemical properties of the region of interest [11]–[14]. While Ramachandran basin allows a holistic description of conformation, this approach lacks statistical description with a focus on the torsion angle distributions of specific sequence and thus in consideration of the circular nature of angles, traditional parametric or non-parametric density estimation methods cannot work properly to approximate Ramachandran distributions; this is also supported by the findings of the current study. As a result, all of this calls for a more unified approach in analysing local protein regions and extracting the high information content from dihedral angle distribution. By extension, capturing sequence information content can facilitate current efforts to build improved predictive models for dihedral angle prediction, protein three-dimensional structure determination and target evaluation for drug screens [13], [14]. To achieve this an additional parameter, the oscillation amplitudes of the atoms around their equilibrium positions (B-factors) in the crystal structures, was used; this relationship is described in the first equation.

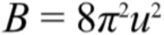

**Equation 1**. B-factor evaluation where oscillation amplitude is u.

While B-factors are used in the atomic form factor calculation to measure scattering amplitude (eq. 2), B-factors have a much more complex influence on atoms and overall structure because of their dependence on conformational disorder, dynamic changes of the sequence seen via the changes in the positional dispersion of B-factors [7], [15].

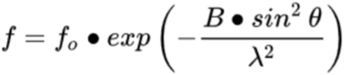

**Equation 2**. Scattering amplitude evaluation where B is B-factor, f_o_ - atomic form factor, λ - the X-ray wavelength.

In addition, B-factors provide means to gain insight on many aspects of molecular dynamics, such as thermal motion paths, protein superimposition and predict the rotameric state of amino acids side-chains [16]–[18]. B-factors were shown to be related to protein packing and depend on three-dimensional structure [14], [19]–[22] and there are numerous other studies investigating protein flexibility through B-factors [22]–[25]. Moreover, B-factors allow the capture of differences between crystal packing sites and biologically relevant protein-protein interaction sites [25]. It becomes apparent that B-factors carry a lot of information on both local and distant protein topologies and by incorporating B-factor estimates we include additional information on the local mobility of a Cα atom. This leads to our derived equation (eq. 4) that provides a fingerprint score or fi-score through the cumulative sum of standard deviation normalised dihedral angles and scaled B-factors divided by the amino acid residue number of a selected region of interest. The fingerprint value captures physicochemical qualities of a region of interest dependent on conformation; moreover, by normalising and scaling we can effectively compare regions of different targets.

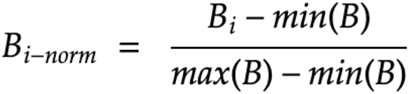

**Equation 3**. Min-max normalisation and scaling of B-factor where B_i-norm_ is scaled B-factor, B_i_ - B-factor for Cα, B_max_ - largest B-factor value for the total protein B-factors for all Cα, B_min_ - smallest B-factor value for the total protein B-factors for all Cα.

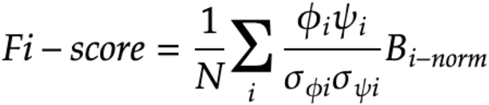

**Equation 4**. Fi-score evaluation where N- is the total number of atoms for which dihedral angle information is available, ϕ and ψ values represent dihedral angles for an Cα atom, σ_ϕ_ and σ_ψ_ represent corresponding standard deviations for the torsion angles and B_i-norm_ is a normalised B-factor value for the Cα atom.

The described methodology could be of great pharmaceutical interest in the identification of families of targets that are affected by drug treatment and to characterise binding sites after mutational studies. For example, when a signalling protein family contains known drug targets, fingerprinting can define additional druggable family members without relying on the scanning of sequence similarity but actually measuring physicochemical parameters [26]. Fi-score visualisation can also aid to accurately evaluate distribution of different regions along the protein sequence by capturing distal information on the interacting amino acids. Furthermore, applying machine learning techniques to cluster fi-scores can reveal dynamically similar sites so that proteins can be scored and grouped prior to computationally expensive *in silico* high through-put screening.

## Methods

### Protein set selection

Structures of proteins were downloaded directly from RCSB Protein Data Bank (PDB) by first selecting proteins based on their features using databases: Pfam 32.0 and Structural Classification of Proteins (SCOP) databases. This diverse set of randomly selected proteins was used for comparative studies (table 1, PDB IDs in bold) which then were analysed using protein BLAST to find good candidates to form protein pairs that showed a varying degree of similarity (table 1, PDB IDs not highlighted). From the initial pool of 200 proteins, only the most reliable 70 structures were then selected from the PDB maintaining diversity of resolution and *R*-factor. All paired proteins were subjected to local alignment to identify regions with as much diversity as possible in their identity and similarity scores. These regions were extracted and sequences were globally aligned to get the final score on the identity, similarity and gaps since only that region of interest will be used for fi-scoring. The alignment and testing was performed with the following tools and parameters MSA (MUSCLE algorithm, default parameters; UGENE software version 1.32), pairwise alignment (Smith-Waterman algorithm-Water (EMBOSS), matrix: BLOSSUM62; gap opening: 10; gap extension: 0.5), global pairwise alignment (Needleman-Wunsch algorithm-Needle (EMBOSS), matrix: BLOSSUM62; gap opening: 10; gap extension: 0.5) and Protein-Blast/ PSI-Blast analyses using default seyngs were employed to assess the sequences.

**Table 1.**
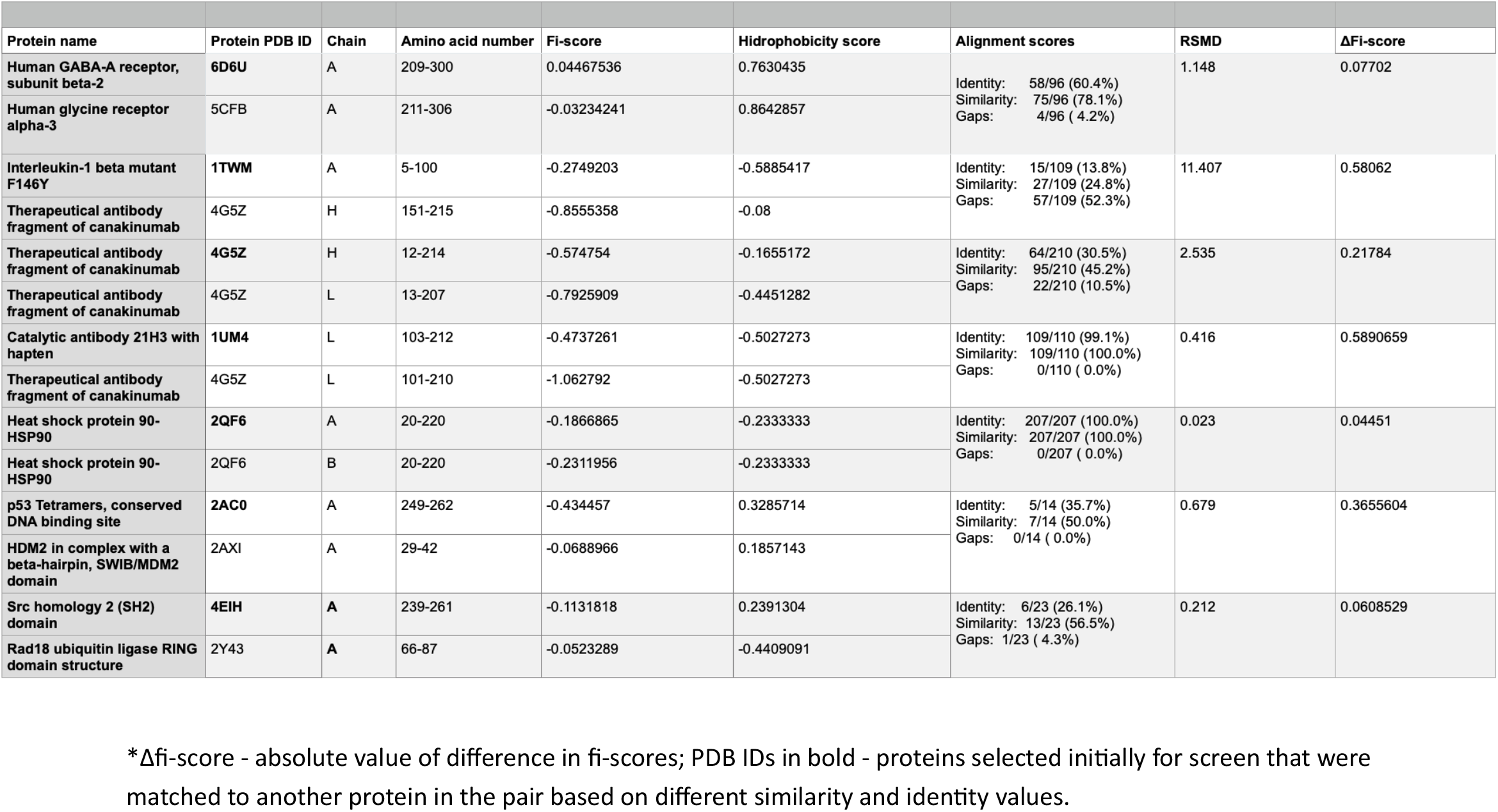
Characterisation and scoring of different target protein regions

| Protein name | Protein PDB ID | Chain | Amino acid number | Fi-score | Hydrophobicity score | Alignment scores | RSMD | ΔFi-score |
| --- | --- | --- | --- | --- | --- | --- | --- | --- |
| Human GABA-A receptor, subunit beta-2 | **6D6U** | A | 209-300 | 0.04467536 | 0.7630435 | Identity: 58/96 (60.4%)<br>Similarity: 75/96 (78.1%)<br>Gaps: 4/96 ( 4.2%) | 1.148 | 0.07702 |
| Human glycine receptor alpha-3 | 5CFB | A | 211-306 | -0.03234241 | 0.8642857 |  |  |  |
| Interleukin-1 beta mutant F146Y | **1TWM** | A | 5-100 | -0.2749203 | -0.5885417 | Identity: 15/109 (13.8%)<br>Similarity: 27/109 (24.8%)<br>Gaps: 57/109 (52.3%) | 11.407 | 0.58062 |
| Therapeutical antibody fragment of canakinumab | 4G5Z | H | 151-215 | -0.8555358 | -0.08 |  |  |  |
| Therapeutical antibody fragment of canakinumab | **4G5Z** | H | 12-214 | -0.574754 | -0.1655172 | Identity: 64/210 (30.5%)<br>Similarity: 95/210 (45.2%)<br>Gaps: 22/210 (10.5%) | 2.535 | 0.21784 |
| Therapeutical antibody fragment of canakinumab | 4G5Z | L | 13-207 | -0.7925909 | -0.4451282 |  |  |  |
| Catalytic antibody 21H3 with hapten | **1UM4** | L | 103-212 | -0.4737261 | -0.5027273 | Identity: 109/110 (99.1%)<br>Similarity: 109/110 (100.0%)<br>Gaps: 0/110 ( 0.0%) | 0.416 | 0.5890659 |
| Therapeutical antibody fragment of canakinumab | 4G5Z | L | 101-210 | -1.062792 | -0.5027273 |  |  |  |
| Heat shock protein 90-HSP90 | **2QF6** | A | 20-220 | -0.1866865 | -0.2333333 | Identity: 207/207 (100.0%)<br>Similarity: 207/207 (100.0%)<br>Gaps: 0/207 ( 0.0%) | 0.023 | 0.04451 |
| Heat shock protein 90-HSP90 | 2QF6 | B | 20-220 | -0.2311956 | -0.2333333 |  |  |  |
| p53 Tetramers, conserved DNA binding site | **2AC0** | A | 249-262 | -0.434457 | 0.3285714 | Identity: 5/14 (35.7%)<br>Similarity: 7/14 (50.0%)<br>Gaps: 0/14 ( 0.0%) | 0.679 | 0.3655604 |
| HDM2 in complex with a beta-hairpin, SWIB/MDM2 domain | 2AXI | A | 29-42 | -0.0688966 | 0.1857143 |  |  |  |
| Src homology 2 (SH2) domain | **4EIH** | A | 239-261 | -0.1131818 | 0.2391304 | Identity: 6/23 (26.1%)<br>Similarity: 13/23 (56.5%)<br>Gaps: 1/23 ( 4.3%) | 0.212 | 0.0608529 |
| Rad18 ubiquitin ligase RING domain structure | 2Y43 | A | 66-87 | -0.0523289 | -0.4409091 |  |  |  |

\*Δfi-score - absolute value of difference in fi-scores; PDB IDs in bold - proteins selected initially for screen that were matched to another protein in the pair based on different similarity and identity values.

### Protein dihedral angle analysis and site scoring

Protein dihedral angles were analysed using R package Bio3D [27] with specific modifications to allow dihedral angle retrieval, fingerprint calculation and visualisation (R studio, version 1.1.463). AddiVonal functionalities were introduced to better capture dihedral angle and B-factor distribution. Hydrophobicity scoring for a selected site was calculated based on Kyte-Doolitle scale.

### Protein visualisa, on and structural analysis

PyMOL (Molecular Graphics System, Version 2.0 Schrödinger, LLC) was used for protein visualisation and superimposition studies as well as structural analysis integrating python code for robust parsing.

### Protein feature capture

Gaussian mixture models (GMMs) (with the following parameters: max_iter=1000, covariance_type=‘full’ or ‘spherical’, tol=0.001, random_state=0)) were implemented to cluster fi-score profiled protein sequences. The number of components for clustering and correction of the over-fiyng was established using Akaike information criterion (AIC) and the Bayesian information criterion (BIC) where the smallest difference between Y_AIC_ and Y_BIC_ information criterion values was used to determine a component number (usually spanning the inflection point of both curves). Python Scikit-Learn GMM (scikit-learn 0.22.2) was used for the above analyses where Gaussian mixture and expectation-maximisation algorithms where defined by the 5-7 equations to estimate the density and distribution of fi-scores for amino acids.

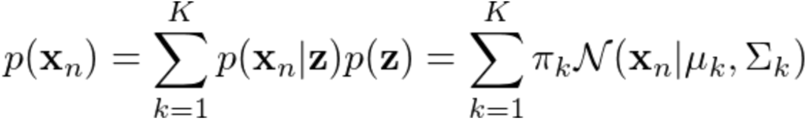

**Equation 5**. Equation defining a Gaussian Mixture; where Σ_k_-covariance for the Gaussian, *K* is the number of clusters of the dataset, μ_k_-cluster centre, π_k_-mixing probability, z - a latent variable defining a probability that data point comes from the Gaussian.

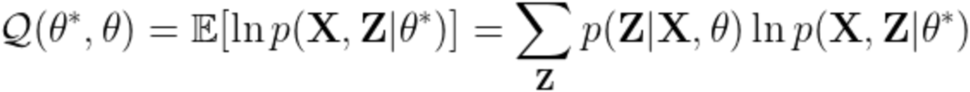

**Equation 6**. Expectation step defining the equation where the current value of the parameters *ϑ* is used* to find the posterior distribution of the latent variables given by P(Z| X,θ*).

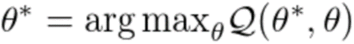

**Equation 7**. Maximisation step defining the equation to find the expectation under the posterior distribution of the latent variables with a new estimate for the parameters.

## Discussion and Results

Protein conformation determination and capturing of the physicochemical properties remains one of the most important topics for drug discovery purposes [28], [29]. That is, defining protein regions that share similar dynamic range is a significant challenge and in order to address that we developed a method to capture side chain as well as mean atomic displacement distribution to provide a value that can aid in comparing and characterising regions of interest which we call a fingerprint score or ‘fi-score’.

Fi-score equation through the use of standard deviation normalised dihedral angles values and scaled B-factor using min-max method allows to effectively compare the resulting score across different targets or sites. We looked into how conformational criteria can be extracted from backbone torsion angles (ψ,ϕ), that follow a very specific local geometry to avoid steric clashes (supl. fig.1); this led us to adopt a standard deviation normalisation for the observed torsional angles. While several different normalisation approaches exist for dihedral angles [30], [31], the mathematical techniques directly depend on the parameter incorporation into further equations which, in our case, needed to be formulated in a way to preserve Ramachandran plot directionality based on positive and negative value so that multiplication operation allowed to predict either β-sheet/strand type of conformation (negative) or α-helix (positive) for the most predominant secondary structure elements (supl. fig.1). When the cumulative score is calculated the ultimate value can indicate the predominance of the said structures and in rarer situations a less dominant conformations, such as a left-handed α-helix. Similarly, B-factor values needed to be scaled since values may be on different scales owing to dissimilar refinement procedures (supl. fig.3) [16], [22], [32], [33]; we applied scaling specifically to take that into account where scale normalised values of B-factors ranged from 0 to 1 allowing them to be conceptually integrated into the fingerprint score equation (supl. fig.2). Finally, dividing the cumulative sum by the number of residues we can measure an average value for the region or fi-score.

A diverse set of randomly selected proteins was used for the comparative studies (table 1) which then allowed us to contrast varied regions of target proteins based on their fi-score, normalised B-factor as well as dihedral angle distribution and sequence alignment data (fig. 1, supl. fig. 2). In addition, the selected region was scored for hydrophobicity and a RMSD value was identified for two target sequences. Target sequences that share higher similarity have closer fi-score values that also correspond to a more similar distribution profile (fig. 2), for example, a sequence from human GABA-A receptor, subunit beta-2 (PDB ID: 6D6U) sharing 78.1% similarity with human glycine receptor alpha-3 (PDB ID: 5CFB) differ by 0.07702 in their fi-scores. When compared to a case of 100% similarity as is for chain A and B of human heat shock protein 90, Δfi-score value drops to 0.04451. However, the sequence similarity alone does not play a defining role as illustrated by catalytic antibody 21H3 (PDB ID: 1UM4) with hapten and therapeutical antibody fragment of canakinumab (PDB ID: 4G5Z) light chains that although share 100% sequence similarity, have almost a half of fi-score difference between them; this is because fi-score captures the 3D distribution of the amino acids, side chain orientation and the predicted atom movements. The slight shifts in the amino acid and their side change orientation (fig. 1) will have a noticeable effect on the fi-score. Moreover, protein regions that have large structural differences as showcased by the interleukin-1 beta mutant F146Y (PDB ID: 1TWM) and therapeutical antibody fragment of canakinumab (PDB ID: 4G5Z) (RMSD=11.407 Å) will have large corresponding differences between fi-score and RMSD values. In many cases smaller Δfi-score values will mean that protein regions have similar structural and physicochemical profiles but in more ambiguous cases hydrophobicity analysis should be included as it indirectly captures the nature of amino acid composition in the selected region as illustrated by the cases (table 1). This information can be especially useful as it allows to compare protein regions of similar mobility or amino acids that have a matching structural profile, for example, the conserved DNA binding site of p53 (PDB ID: 2AC0) is quite similar to a region of SWIB/MDM2 domain (PDB ID: 2AXI) and a similar profile can be seen for heavy and light chains of therapeutic antibody fragment of canakinumab (PDB ID: 4G5Z) (fig.1, table 1). These examples illustrate that depending on 3D organisation of a region of interest, consertiative substitutions of amino acids, dihedral angle and B-factor values will have an impact on individual fi-score values for amino acids and cumulative score. There are many studies that support these findings as it has been established that B-factors can be used to identify flexibility in proteins and can also be linked to hydrophilicity as well as absolute net charge [12], [23], [25], [34]–[37]). B-factors can also aid in identifying biologically active small molecules for a site of interest [24], [25], [36], [38].

**Figure 1.**
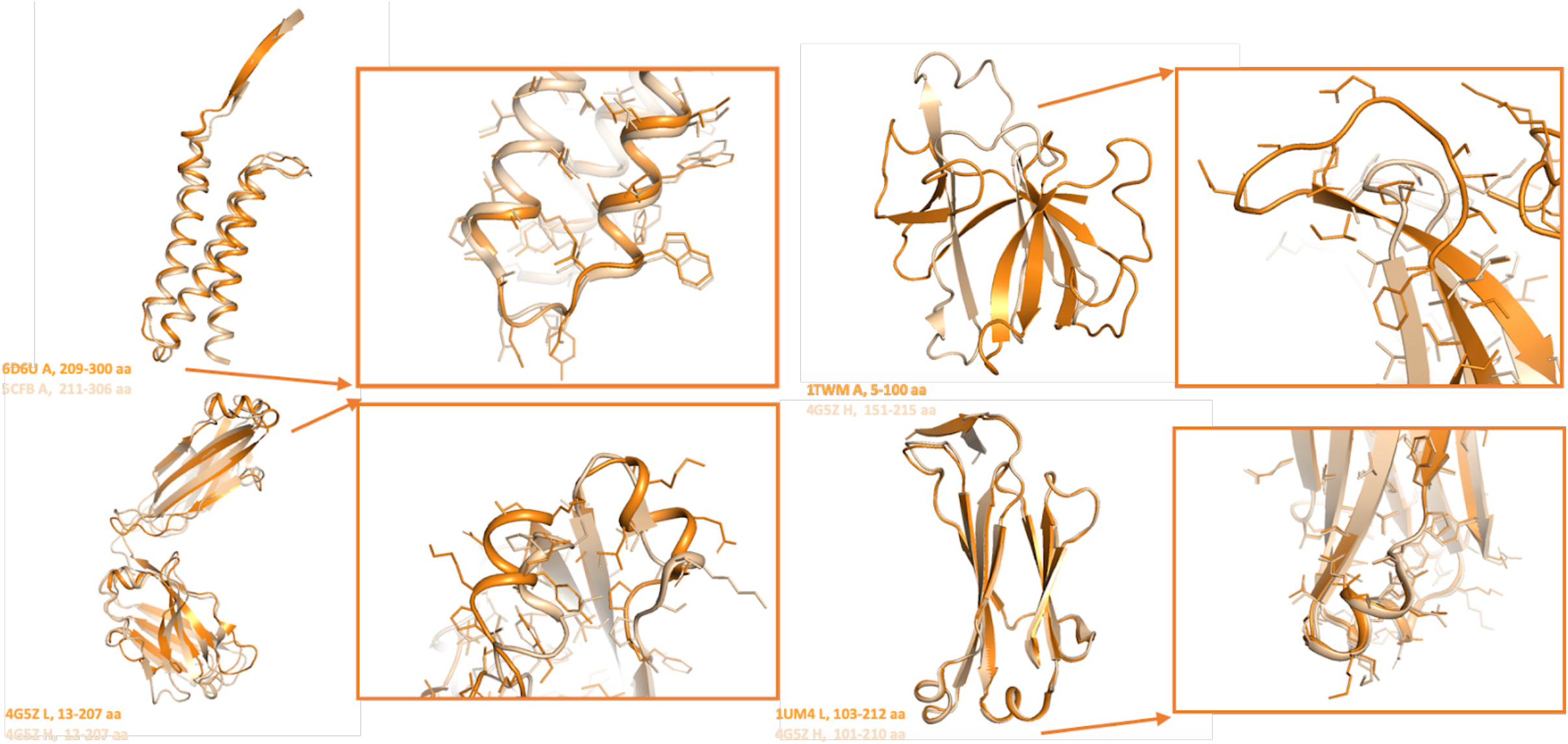
Superimposition of representative examples of human protein regions listed in table 1 where PDB ID and colour is specified next to the structure.

**Figure 2.**
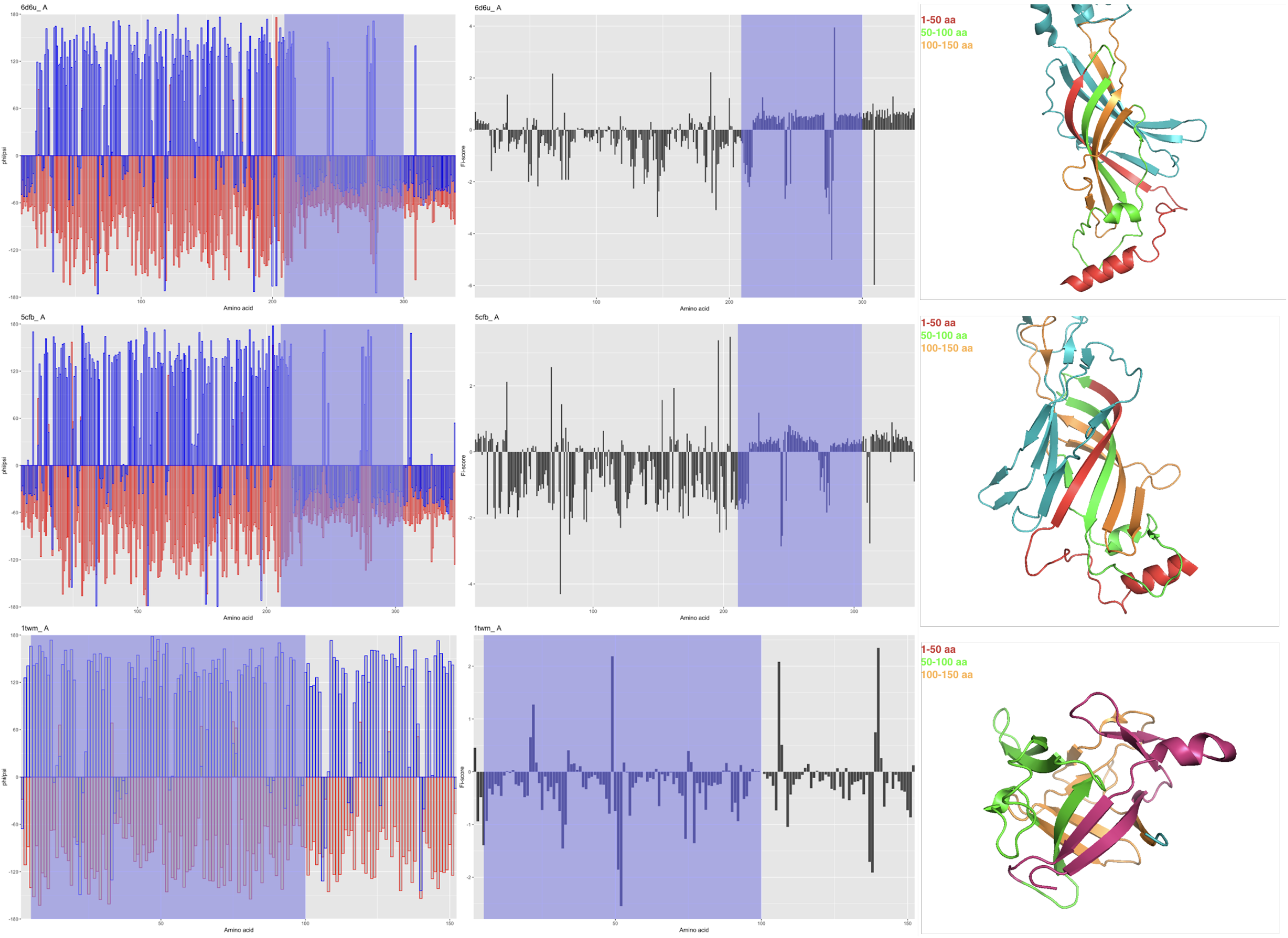
Dihedral angle and fi-score distribution, respectively lef and right panels, where ϕ dihedral angle is represented in red and ψ - blue. Blue region delineates the protein section that was used to calculate fi-score and supporting information as per table 1. Right panels of the corresponding protein structure are colour coded in arbitrary increments of 50 amino acids.

Another important criterion is the region size selected for the analysis since fi-score encapsulates conformational information and does not rely on sequence values alone (where sequence influence arises in a form of dihedral angle and B- factor distribution) [19], [29], [33], [34], [39].

Selected window size for the analysis will have an effect on what information is contained within the fi-score; a smaller window size of approximately 20-50 amino acids can reflect the profile of an average motif in a protein (fig. 2), however, larger window sizes averaging 100 or more amino acids reveal averaged physicochemical information of that larger window size (fig. 1). This is especially evident when looking at individual fi-score values per amino acid (fig. 2) where fi-score distribution captures different protein regions not easily recognised by looking at dihedral angle or B- factor distribution alone (fig. 2, supl. fig. 1&2).

These obsertiations prompted us to investigate a varied set of proteins with various structural elements ranging from β-sheets/strands to α-helices as well as mixed or disordered regions (table 2, fig. 3). By narrowing down to a unique structural motif we can capture not only its physicochemical properties but also reliably categorise it to either α-helix or β-sheets/strands-like structures. However, structures that are a mixture of several components, e.g., 6S34 (table 2), might have opposite sign values or values closer to 0 for fi-scores because some less predominant structures of α-helices and β-sheets occupy negative and positive basins, respectively.

**Table 2.** Structural motif and domain physicochemical characterisation for selected protein structural elements and motifs

| Protein PDB ID | Description | Amino acid number | Fi-score | Hidrophobicity score |
| --- | --- | --- | --- | --- |
| 6S34 | α-helix with disordered linker | 2-19 | -0.1727632 | 0.3277778 |
| 6S34 | Iregular C-end α-helix | 11-17 | 0.6483593 | -0.7 |
| 6RZ3, A | Single outer α-helix of cellular tumour antigen p53 | 278-290 | 0.06747168 | -2.646154 |
| 6RZ3, B | Outer α-helix surrounded by smaller helices and unstructured regions of the carboxyl-terminal conserved region of inhibitor of apoptosis-stimulating protein of p53 (IASPP) | 741-752 | 0.1493196 | 0.175 |
| 6RZ3, B | Four α-helices with long stretches of disordered linkers of the carboxyl-terminal conserved region of inhibitor of apoptosis-stimulating protein of p53 (IASPP) | 661-715 | -0.02906267 | 0.4854545 |
| 1W7B | Contorted α-helix joining two separate α-helices | 154-176 | -0.4146927 | -0.1521739 |
| 1W7B | Contorted α-helix joining three separate α-helices | 238-276 | 0.02634468 | -0.4 |
| 1W7B | Single outer α-helix | 187-200 | 0.0995633 | -1.264286 |
| 1W7B | Single region of left handed α-helix like structure | 89-93 | -1.233356 | 0.72 |
| 1DEE, G | Protein A ImG binding domain, α-helical motif, chain G | 1810-1852 | 0.122619 | -0.7627907 |
| 1QCF | Antiparallel β-strands | 183-207 | -0.8710873 | -1.108 |
| 1QCF | Antiparallel β-strands | 266-296 | -0.2612031 | -0.4129032 |
| 1A0S, R | β-barrel, with three random coil motifs | 76-476 | -0.3455974 | -0.4897756 |
| 1A0S, R | Three antiparallel β-sheets; outer pore region | 351-401 | -0.3455974 | -0.4960784 |
| 1A0S, R | Three antiparallel β-sheets connected via α-helix; inner pore region | 91-146 | -0.2178924 | -0.5375 |
| 8TIM | TIM barrel | 11-241 | -0.04860643 | -0.1311688 |
| 8TIM | TIM barrel motif of two β-sheets and α-helix | 6-41 | -0.1594032 | -0.09166667 |
| 1DEE, E | ImG Fab β-sheet, chain E | 2006-2076 | -0.6376855 | -0.2239437 |
| 1DEE, A | ImG Fab β-sheet, chain A | 6-76 | -0.2326242 | -0.2239437 |
| 6RZ3, B | A single stretch of β-sheet strand | 806-811 | -1.700738 | -1.016667 |
| 6RZ3, B | Disordered region of a protein | 764-781 | -0.7002029 | -0.5944444 |

**Figure 3.**
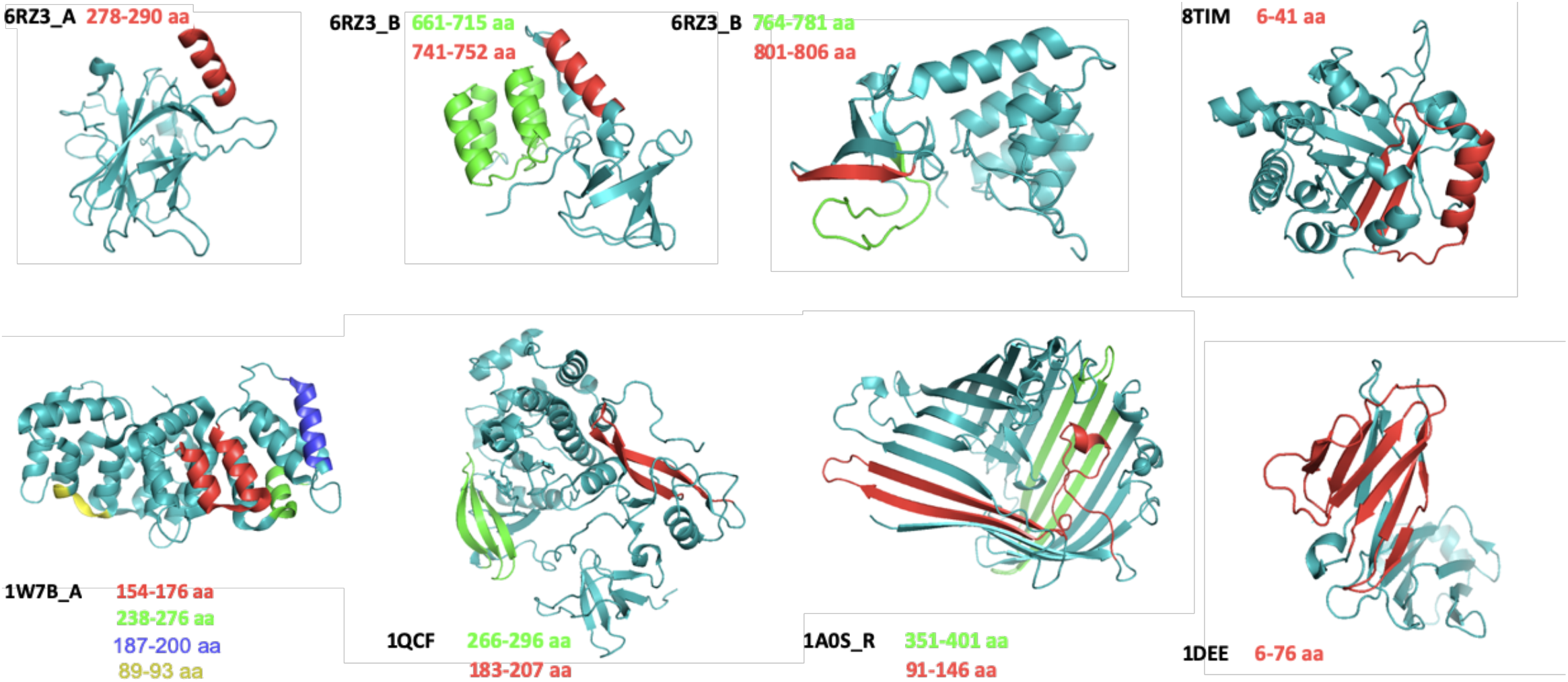
Representative protein structural motifs and regions from table 2 are color coded for specific amino acids.

The higher absolute fi-score value the more flexible region is likely to be, for example, the outer α-helix of DNA-binding domain of p53 (chain A, PDB: 6RZ34_A; table 2, colour - red) might appear untethered and relatively flexible; however, based on low absolute fi-score value and further inspection on inter-chain H-bond formation (supl. fig. 3) this structure is accurately predicted to be of limited dynamic range. Other α-helices complexes within the chain B of the inhibitor of apoptosis-stimulating protein of p53 (iASPP; PDB: 6RZ34_B; table 2) are of varying flexibility because they are at the contact point between two chains, specifically: a shorter α- helix (chain B) is less constrained by polar contacts and interaction surface then the other participating elements of this contact site. Similar patterns can be observed in α-helices and their complexes of annexin A2 (PDB: 1W7B, table 2, figure 3) where flexibility and fi-score value depends on the conformation. In the case of β-strands and β-sheets, these secondary structure elements, have the same trend of higher flexibility with higher fi-score values (iASPP; PDB: 6RZ34, chain B; table 2, figure 3). More compact sites have minimal space of side chain and motif movement as can be seen in triose phosphate isomerase (ODB: 8TIM, chain A; table 2, figure 3). Finally, disordered regions or regions that combine several secondary structure elements might have sign value depending on the dominating structural motif. All of the above findings are supported by earlier studies showing that B-factors can act as indicators of the relative vibrational motion of atoms where low values belong to a well-ordered site, and the highest values come from the most flexible regions [10], [15], [33], [38], [40].

Based on the findings that fi-score captures and allows to differentiate among different protein regions, we wanted to check if applying clustering would allow us to categorise fi-score values as we have already observed clear distribution patterns (fig.1). However, some protein regions might be in transition states and thus, have similar or overlapping fi-score values and in order to address that we selected Gaussian mixture models (GMM) [9], [22], [41], [42]. GMM is often categorised as a clustering algorithm, but it has much broader implications functioning as a density estimator. Since fundamentally GMM is a generative probabilistic model, this algorithm was chosen to describe the distribution of the fi-scores.

The covariance type for the fits of the majority of studied cases was left to be modelled as an ellipse of an arbitrary orientation for each cluster and the optimal number of components for a given dataset was determined using AIC and BIC approaches to avoid overfiyng. Fi-score clustering revealed that GMM allows not only to capture different secondary structure elements (fig. 4) but at the same time group them into physicohemically similar units based on dihedral angle determined side chain orientation and B-factor predicted amino acid oscillations amplitude. In the case study of catalytic antibody 21H3 with hapten (PDB: 1UM4, chains H and L; fig. 4, sup. fig.5) we can see that fi-score evaluation and clustering successfully determined complementarity determining region β-turns and different β-strands in immunoglobulin fold. The heavy and light chain contact sites are also captured through different chain topology and relevant atomic movement. Another case study of triose phosphate isomerase (PDB: 8TIM) demonstrated how fi-score based method allows to differentiate secondary structure elements of α-helices and loops at the C-terminal ends of the β-barrel which are known to be involved in catalytic activity [43]. Similarly, N-terminal loops performing a stabilising function were also distinguished from the surrounding structural elements. This ubiquitous enzyme fold can be further resolved into different motifs of interchanging α-helices and β-strands forming the structure’s core (fig. 4).

**Figure 4.**
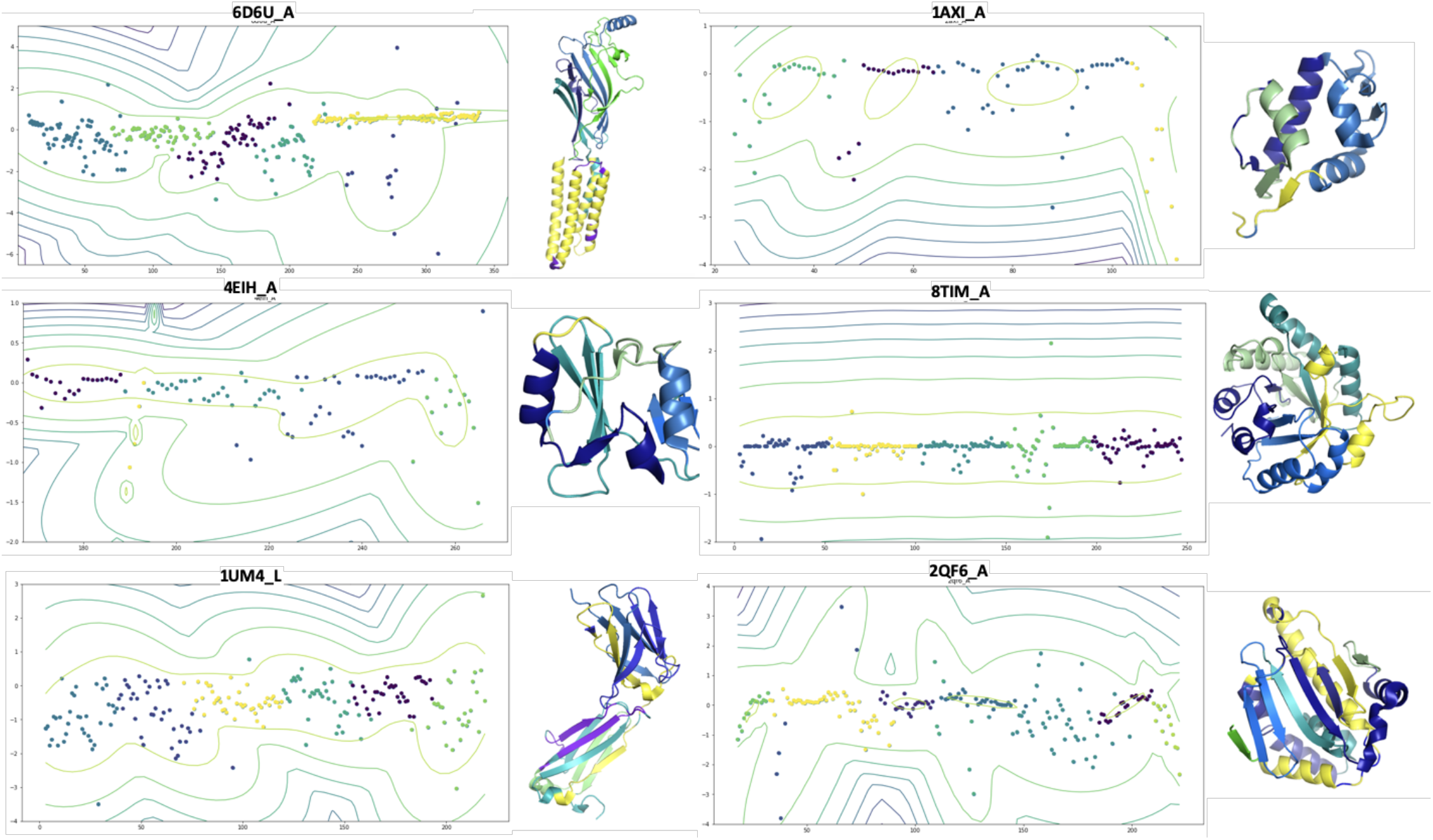
Representative proteins and their fi-score clustering based on density estimation GMM where colours of the clusters as well as density lines match structural element colours of a protein on the right.

As illustrated, fi-score entered analysis can be a powerful tool to gain insight into structural topology of a target of interest. Furthermore, by including density estimation contours we can predict the changes of a protein region if, for example, the structure is not in a crystal but in a solution [44], [45]. This method could also be expanded to estimate effects of mutations and what changes in fi-score value are the most optimal. Finally, drug screening studies can benefit from classifying target sites and cross-referencing with known binders which could reduce off-target effects as well as allow to address and better understand cases of unspecific binding or dynamic instability.

## Conclusion

We provide a new method to characterise proteins prior to *in silico* screening by evaluating potentially dynamically active regions or predicting sites that share similar qualities in side chain distribution and movement. Incorporating fi-score with other physicochemical parameters, such as hydrophobicity, could greatly improve detecting valuable multiple sites within a target or capturing similar profiles across different targets which in turn could be subjected to docking studies. Moreover, we showed that by using machine learning approaches we can expand and speed-up the analysis of multiple targets by extracting and defining structural elements and motifs of various proteins. Fi-score focused analysis can aid in not only primary target selection studies but also advance drug or biologics formulation methods by evaluating potential binding sites or interaction surfaces. In summary, this innovative biophysical analysis method could significantly improve target selection, pre-screening analysis and speed-up biologics engineering.

## ABBREVIATIONS

HTS: high-throughput screening
PLI: target protein–ligand interactions
AIC: Akaike information criterion
BIC: Bayesian information criterion

## ACKNOLEGEMENTS

The authors would like to thank Dr. Farhad Forouhar, Columbia University, for kindly offering technical advice on structure visualisation

