## Supplementary material for "Fi-score: a novel approach to characterise protein topology and aid in drug discovery studies": suplementary_materials

### Supplementary materials

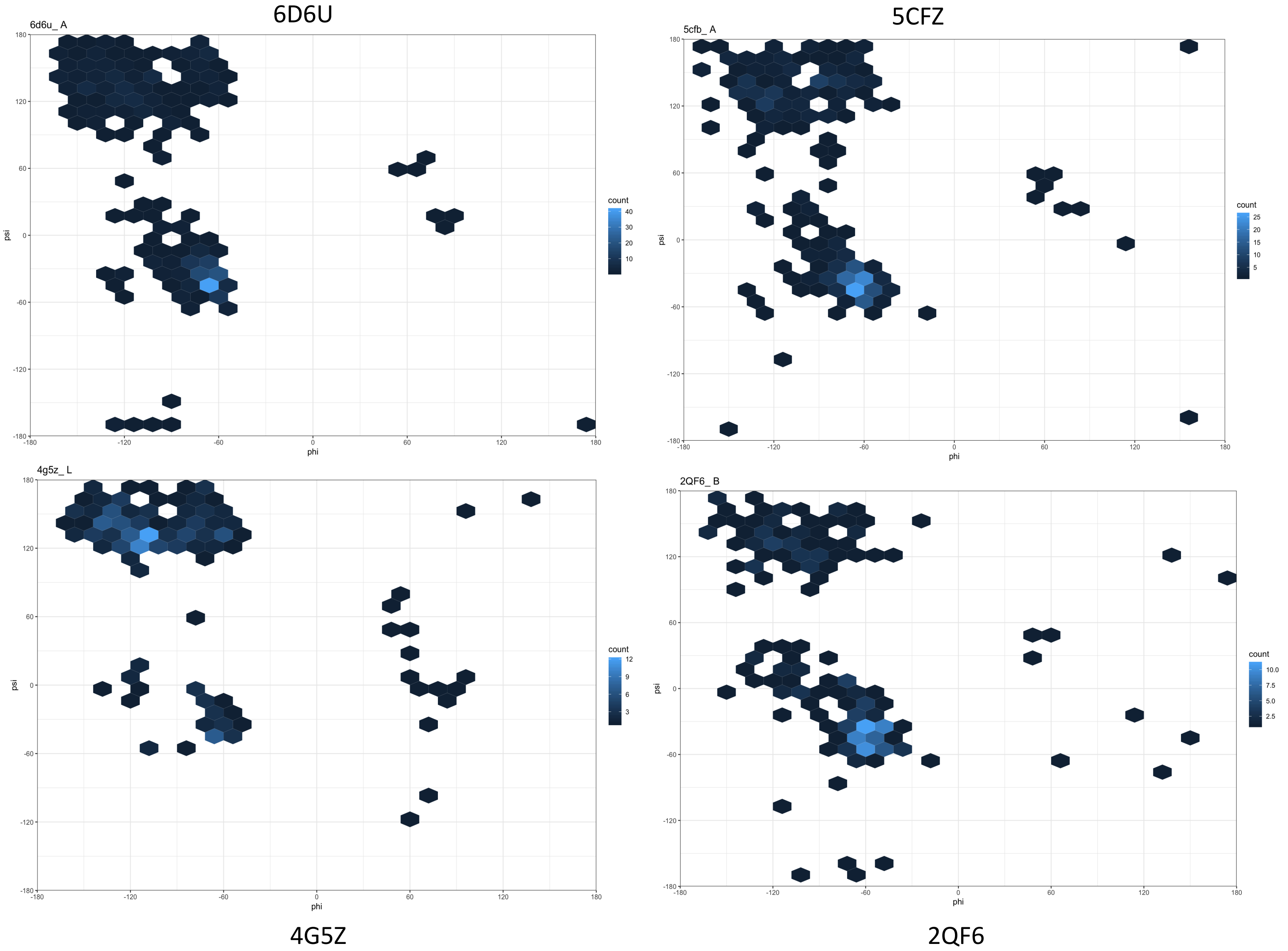


**Supplementary figure 1**. Representative examples of Ramachandran plots for representative proteins PDB: 6D6U(A chain), 5CFZ (A chain), 4G5Z (L chain), 2QF6 (chain B).

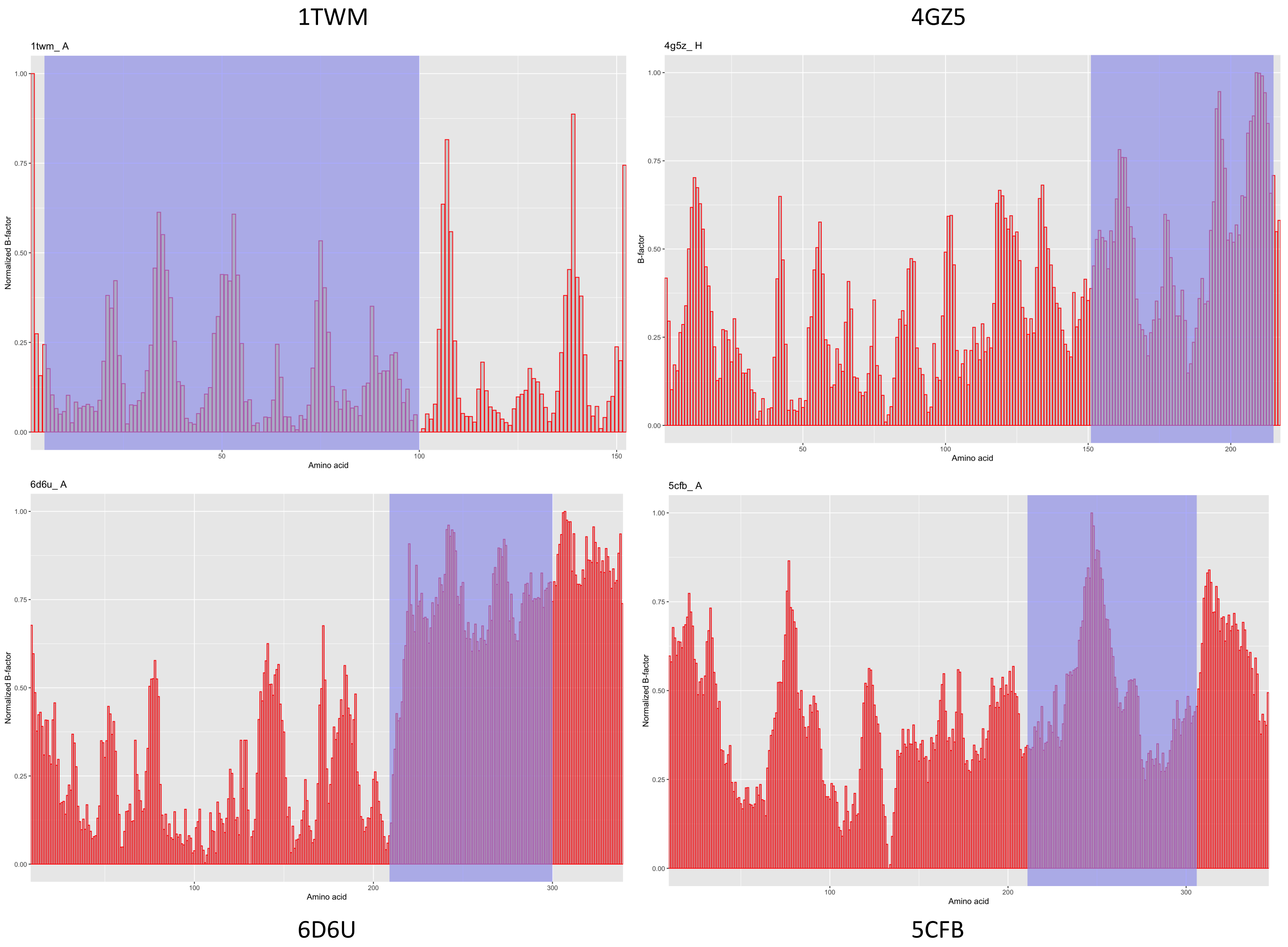


**Supplementary figure 2**. Representative examples of scaled B-factor value distribution (from 0 to 1) for proteins PDB ID: 1TWM (chain A), 4G5Z (H chain),6D6U (A chain), 5CFZ (A chain), where shaded blue region represents analyzed regions (table 1).

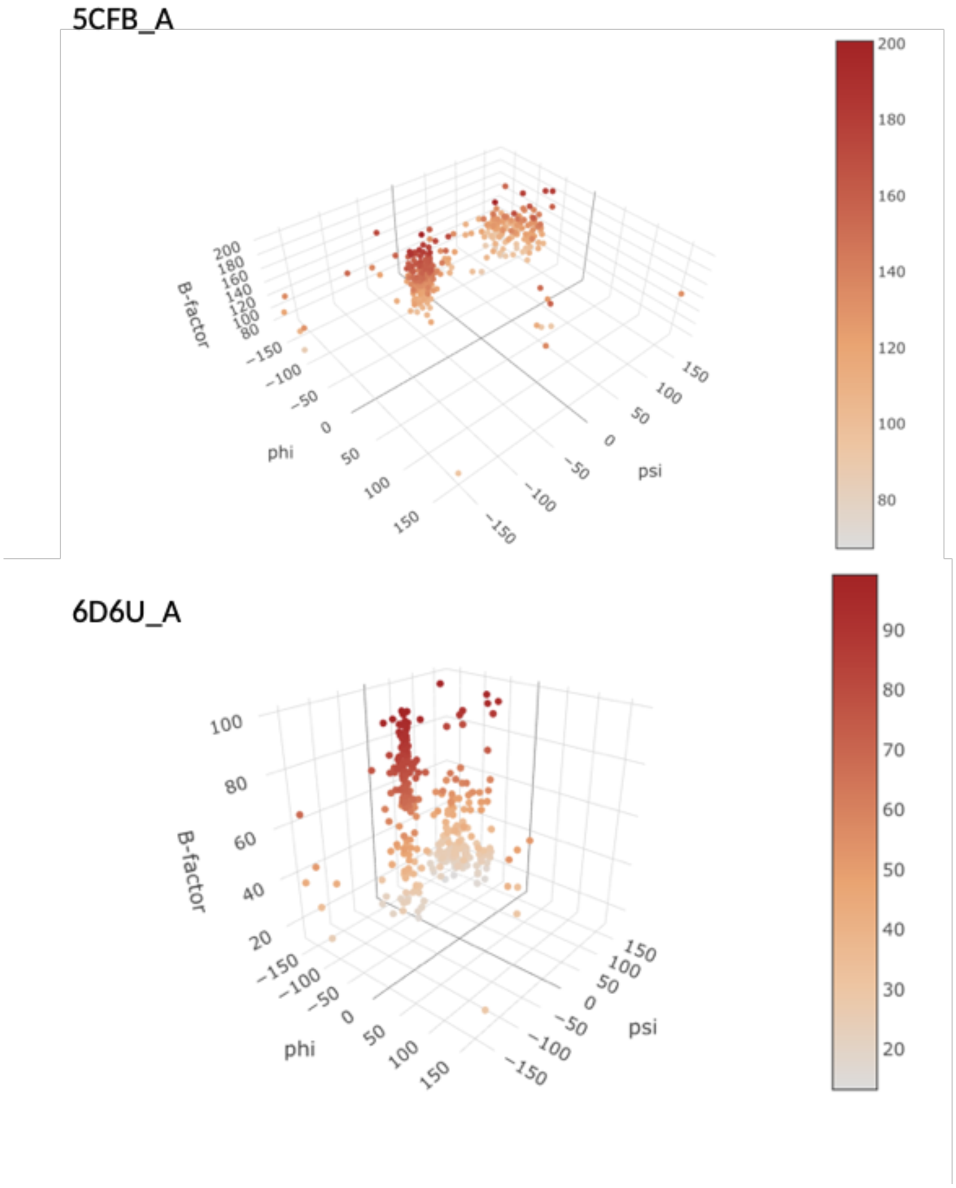


**Supplementary figure 3**. Representative examples of torsion angle and B-factor value distribution for human GABA-A receptor, subunit beta-2 (PDB: 6D6U, chain A) and human glycine receptor alpha-3 (PDB: 5CFB, chain A) where a scale represents B-factor value without normalization.

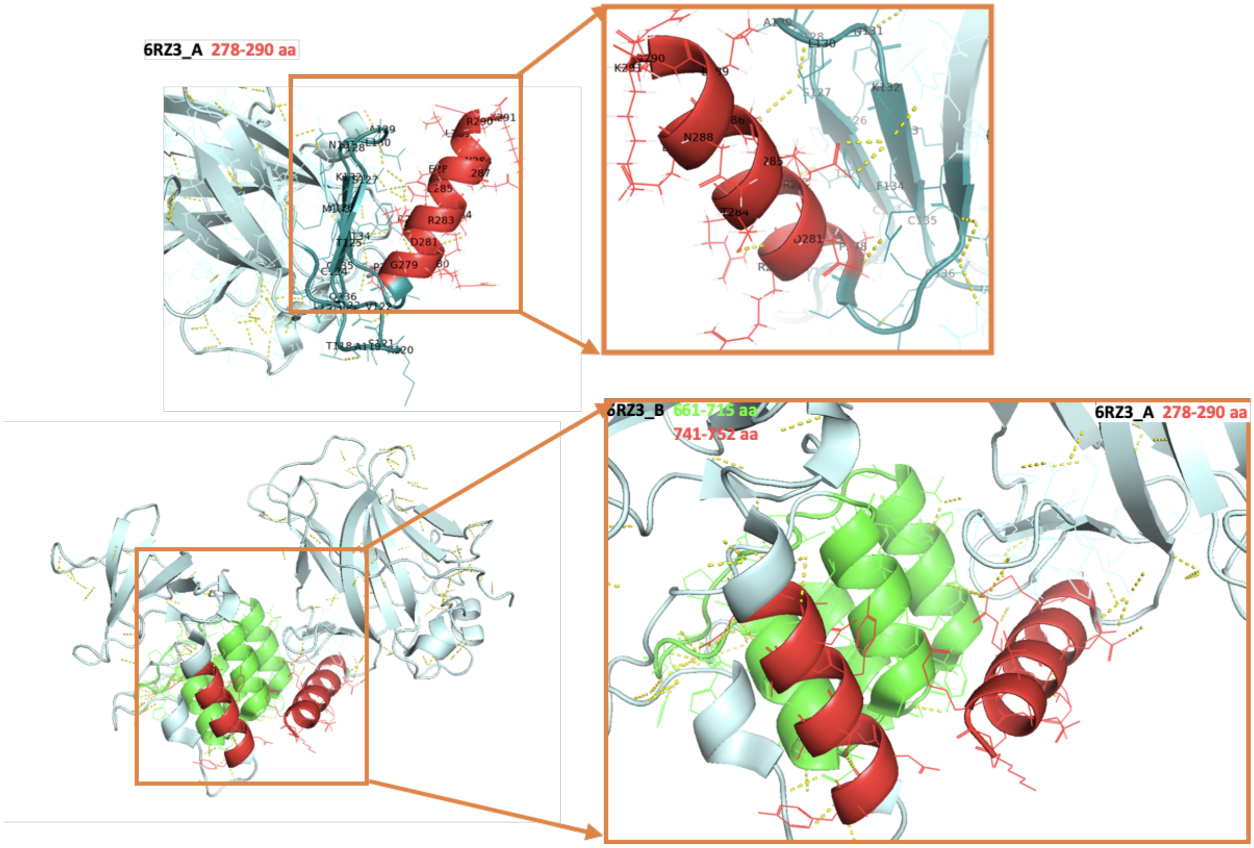


**Supplementary figure 4**. Region of a single outer α-helix of cellular tumor antigen p53 (PDB: 6RZ3, chain A) (top panel) and a contact site between the outer α-helix of cellular tumor antigen p53 (PDB: 6RZ3, chain A) and the carboxyl-terminal conserved region of inhibitor of apoptosis-stimulating protein of p53 (iASPP) (PDB: 6RZ3, chain B) where yellow dotted lines represent interchain polar contacts (bottom panel).

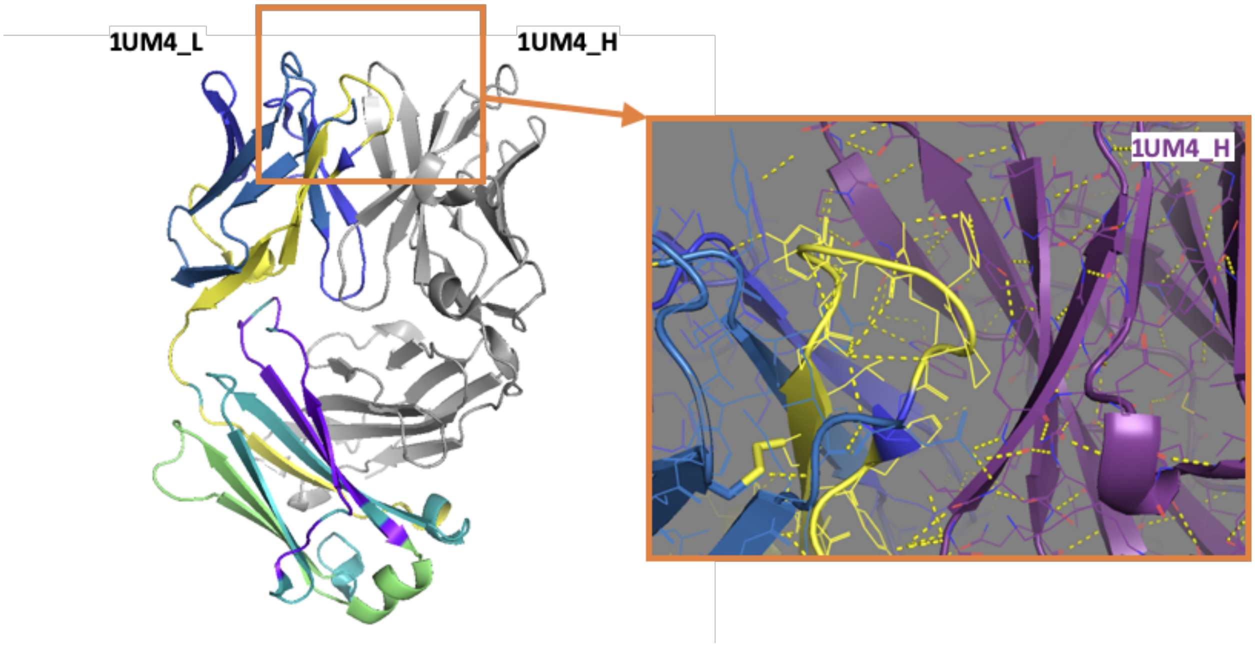


**Supplementary figure 5**. Catalytic antibody 21H3 with hapten (PDB: 1UM4, chain H and L) where N-terminal heavy and light chain contact site are shown in a close-up with polar contacts depicted in a dashed yellow line.
